# Spike sorting with Gaussian mixture models

**DOI:** 10.1101/248864

**Authors:** Bryan C. Souza, Vítor Lopes-dos-Santos, João Bacelo, Adriano B. L. Tort

## Abstract

The shape of extracellularly recorded action potentials is a product of several variables, such as the biophysical and anatomical properties of the neuron and the relative position of the electrode. This allows for isolating spikes of different neurons recorded in the same channel into clusters based on waveform features. However, correctly classifying spike waveforms into their underlying neuronal sources remains a main challenge. This process, called spike sorting, typically consists of two steps: (1) extracting relevant waveform features (e.g., height, width), and (2) clustering them into non-overlapping groups believed to correspond to different neurons. In this study, we explored the performance of Gaussian mixture models (GMMs) in these two steps. We extracted relevant waveform features using a combination of common techniques (e.g., principal components and wavelets) and GMM fitting parameters (e.g., standard deviations and peak distances). Then, we developed an approach to perform unsupervised clustering using GMMs, which estimates cluster properties in a data-driven way. Our results show that the proposed GMM-based framework outperforms previously established methods when using realistic simulations of extracellular spikes and actual extracellular recordings to evaluate sorting performance. We also discuss potentially better techniques for feature extraction than the widely used principal components. Finally, we provide a friendly graphical user interface in MATLAB to run our algorithm, which allows for manual adjustment of the automatic results.

## Introduction

Analyzing the activity of neurons recorded with extracellular electrodes is one of the main techniques to study the brain in vivo. To that end, it is necessary to reliably identify spikes of different neurons recorded from the same electrode. In principle, this can be partially achieved because the extracellular waveform of action potentials varies depending on biophysical and morphological properties of the cells, as well as on the relative position of the electrode [1-3]. Thus, the detected spikes can be assigned into clusters of similar waveforms that correspond to different neurons. This clustering procedure is also known as ‘spike sorting’ and constitutes a crucial step before spike train analysis.

Many algorithms have been developed to deal with this problem [4-6]. One of the main challenges is to automatically identify which features of the spike waveform should be used for classification. In fact, as important as the clustering algorithm per se is the preceding processing step, often referred to as dimensionality reduction [7]. This step aims at transforming the set of waveforms into a small representation in order to reduce noise and provide the most informative components to be used by the clustering algorithm.

A largely used technique for dimensionality reduction is the principal component analysis (PCA) [4,8,9]. Representing the data in terms of a principal component (PC) means that each waveform is redefined as a weighted sum of its values, a linear combination determined by the PC weights; this transforms the entire waveform into a single number. By definition, the first PC is the axis (or linear combination) which captures most variance of the data. The second PC is the axis that captures most variance orthogonal to the first PC, and so on. In this way, waveforms might be represented by a small set of uncorrelated (orthogonal) components that capture most of their variance.

Although it is convenient to represent the data in a set of uncorrelated components, the main caveat of PCA-based sorting is the core assumption that feature variance would be proportional to its capacity of isolating neurons (clustering separability), which may not be the case. Alternatively, one can rescale the data in order to bias PCA to primarily extract multimodal components relevant for clustering, a technique called weighted-PCA (wPCA) [10-12]. In this framework, each time point (sample) of the waveform is divided by its variance (across waveforms) and multiplied by an estimate of its clustering separability. The challenge, then, is to find the optimal method to make such estimate in an unsupervised manner.

A common alternative to PCA is the wavelet decomposition (WD) [13-16], which provides a time-frequency representation of the signal. The WD transforms each waveform into a set of wavelet coefficients, which isolate frequency (or time scale) components at a certain location in time. Similarly to PCA, one can then obtain a small set of wavelet coefficients that capture localized frequency components relevant for spike classification, thus reducing dimensionality. As in the wPCA, estimates of clustering separability are required to select a mixture of relevant components (see Materials and Methods).

Once the complexity of the data is reduced, the next step is to perform clustering with the selected features. Manual selection of each cluster with the aid of visualization tools is a difficult task when there are too many neurons, besides introducing human bias. To automatize this step, previous approaches have used unsupervised clustering based on template matching [17], the distance between cluster centers (i.e., k-means) [4,13], or statistical models of the data (see references [4,11,18-21]).

In this work, we propose a spike sorting framework using Gaussian mixture models (GMMs), a statistical model that fits the data using a mixture of Gaussian distributions. We combined GMMs with different feature extraction techniques (PCA, wPCA and WD) and explored the multiple ways of reducing dimensionality to relevant features. Additionally, GMMs were also used to estimate the number of clusters and classify spikes. We tested our approach using datasets with simulated and actual waveforms and compared its performance with two other mixture-models-based spike sorting algorithms – EToS [10,11] and KlustaKwik [8,9]. Our method showed similar or better performance concerning benchmark attributes such as signal-to-noise ratio, stability and symmetry. We finish by presenting a MATLAB graphical user interface to perform spike sorting based on GMMs, which we made freely available online.

## Materials and Methods

### Theoretical background

#### Principal components analysis

Principal components analysis (PCA) is a linear transformation that maps the data **X** into a new orthogonal basis maximizing the variance of each dimension (see Fig 1B). It is formally defined by **Y** = **XA**, where **X** is an *s*-by-*n* data matrix with *s* observations (i.e., number of spikes) of *n* dimensions (i.e., number of time samples from each waveform), and **A** is an *n*-by-*n* matrix whose columns are formed by the eigenvectors of the covariance matrix of **X**, which are called principal components (PCs). Each column of **Y** corresponds to the scores of a PC, that is, the projection of an **X** column over the direction defined by the PC. The eigenvalue associated to the eigenvector defining the PC denotes the variance of the data in that direction. Notice that projecting the data in a PC direction is equivalent to performing a linear combination of the *n* dimensions of **X**. The PCs are usually sorted in a decreasing order of eigenvalues so that the scores of the first PCs (columns of **Y**) carry most of the variance of the data, and are thus used for further analyses (dimensionality reduction).

**Fig. 1.**
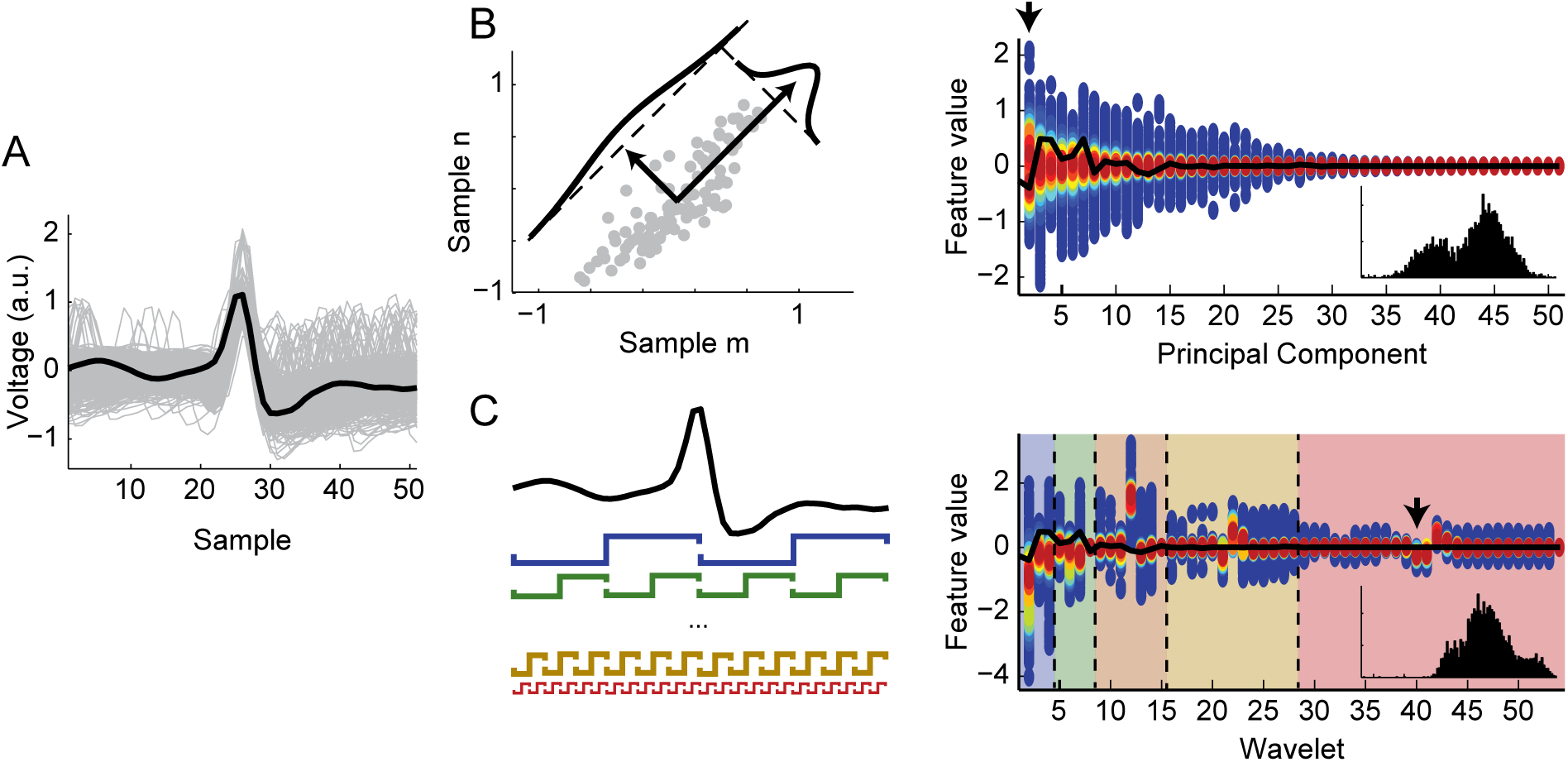
Schematic representation of wavelet decomposition and principal component analysis. **A.** Detected spike waveforms (gray traces) and a particular example (black line). **B.** Principal component analysis of the waveforms. (Left) Scatter plot of two sample points of the waveforms. The principal components (PCs) are successively defined as orthogonal directions maximizing the data variance (black arrows in this example). (Right) Color-coded histogram of the PC scores extracted from the waveforms in A. Warmer colors code for a higher density of waveforms with the same score value. Black line shows the PC scores of the highlighted example in A. Inset shows the histogram of the first PC scores (black arrow). **C.** (Left) Haar wavelet decomposition. Each waveform was decomposed using a set of Haar wavelets of multiple scales and non-overlapping translations (colored functions). (Right) Color-coded histogram of the wavelet coefficients separated by different scales (background colors). Black line shows the wavelet coefficients of the highlighted example in A. Black arrow indicates the wavelet coefficients whose distribution is shown in the inset.

A related procedure, called weighted-PCA (wPCA) [11,12], relies on first normalizing the variance of each dimension n before performing the PCA. This can be used to give more weight to the dimensions that are deemed more relevant (S1 Fig).

#### Wavelet decomposition

A discrete wavelet transform is a time-frequency decomposition that consists in computing the inner product of each waveform *i* (a row of **X**, denoted as x_i_[n]) with scaled and translated versions of the mother wavelet function. This wavelet decomposition (WD) is formally defined as:

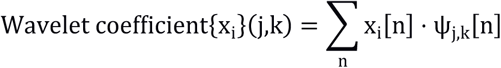

with

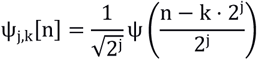

where ψ is the mother wavelet and *j* and *k* are integers representing the scaling and translation factors, respectively. Convolving the signal with the wavelet of a given scale can be seen as a filtering procedure, with the wavelet as a kernel. Here the WD was computed using the multi-resolution approach based on Haar wavelets [22] (Fig 1C). Briefly, this approach uses a Haar wavelet to filter the signal (high-pass filter) and downsamples it by 2 to obtain the wavelet coefficients of the fast scale. The original signal is also filtered using a complementary kernel (a quadrature mirror filter of the wavelet), so that it separates the low-frequency components of the signal (low-pass filter). The procedure is successively repeated using as input the low-frequency signal downsampled by 2, until the desired level of filtering (4 in our case). Because of the downsampling, the next level of filtering captures wavelet coefficients in a slower scale.

#### Gaussian mixture models

A Gaussian mixture model (GMM) is a probabilistic model that assumes the data comes from a combination of *k* Gaussians distributions, with independent means, variances and weights (amplitudes). The GMM for an empirical distribution of feature values can be formally defined as:

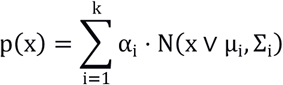

where μ_i_ and Σ_i_, are the mean and covariance matrix of the ith Gaussian, respectively, and α_i_, is the probability that *x* belongs to the ith Gaussian, or the Gaussian weight.

Except for the number of Gaussians, the parameters of the model are iteratively defined in a data-driven manner. We estimated these three parameters θ (mean, covariance and Gaussian weight) using an expectation-maximization (EM) algorithm that searches for the maximum a posteriori probability [23]. Briefly, the EM consists in two steps that are iterated until convergence. In the first step the expected value of *p*(*x*,*z*) - where *z* is the (unknown) membership of the observation *x* - is estimated given an initial set of θ. The second step consists in updating the parameters so that it maximizes the probability of *x* in the model.

Because the resulting GMM depends on the initial conditions, we computed 10 replicates for each fitting and kept the ones with highest log likelihood. We set the stopping criterion as 10^4^ interactions or a convergence threshold (10^-6^ percentage of change in *log p(x)*). In a later step of our algorithm (see below), we used a modified version of the EM in which we could set the Gaussian centers of the model. In this version, which we refer to as “fixed-mean GMM”, only the α and Σ parameters were updated in the first step so that the initial and final μ were the same.

### GMM-based spike sorting

Our proposed algorithm can be separated in two main steps: feature extraction and clustering. Each step uses GMMs, as detailed below:

#### Feature extraction

For feature extraction, we performed either PCA or WD on the set of waveforms (Fig 1). Once the waveforms are transformed into such features (PC scores or wavelet coefficients), a subset of them are selected for clustering. This selection is crucial since the quality of clustering is highly dependent on the amount of spike identity information conveyed by the features. We investigated selection criteria based on GMM, which we explain next.

The probability function for values of a given feature (PC score or wavelet coefficient) was estimated by fitting a GMM with eight Gaussian functions to its empirical distribution (discretized with 100 equally spaced bins across the data range). We computed four different metrics of clustering separability. Three of them were based on GMM, which depended on the (1) peaks and (2) inflection points of the mixture, as well as on the (3) distance between the individual Gaussians (Fig 2). The peak-(I_peak_) and inflection-based (I_inf_) metrics were defined as:

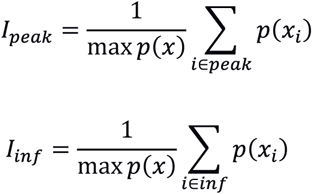

where p(x_i_) is the probability of the model at either a peak or an inflection point x_i_ (see Fig 2B). Intuitively, these metrics are the sum of the model probability at the peaks or inflection points normalized by the highest probability. The third metric, referred to as the distance metric (I_dist_), was computed from the normalized distance between each pair of Gaussians:

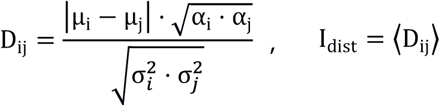

where μ, σ and α are, respectively, the mean, standard deviation and weight of Gaussians *i* and *j* (Fig 2C), and the brackets denote the median over all *i* and *j*. In other words, *D*_*ij*_ measures how distant two Gaussians are after correcting them by their standard deviation. Because the amplitude of each Gaussian can influence the amount of data they separate, we also weighted them by their amplitude, so that a Gaussian pair with high amplitude is more distant than one with low amplitude.

**Fig. 2.**
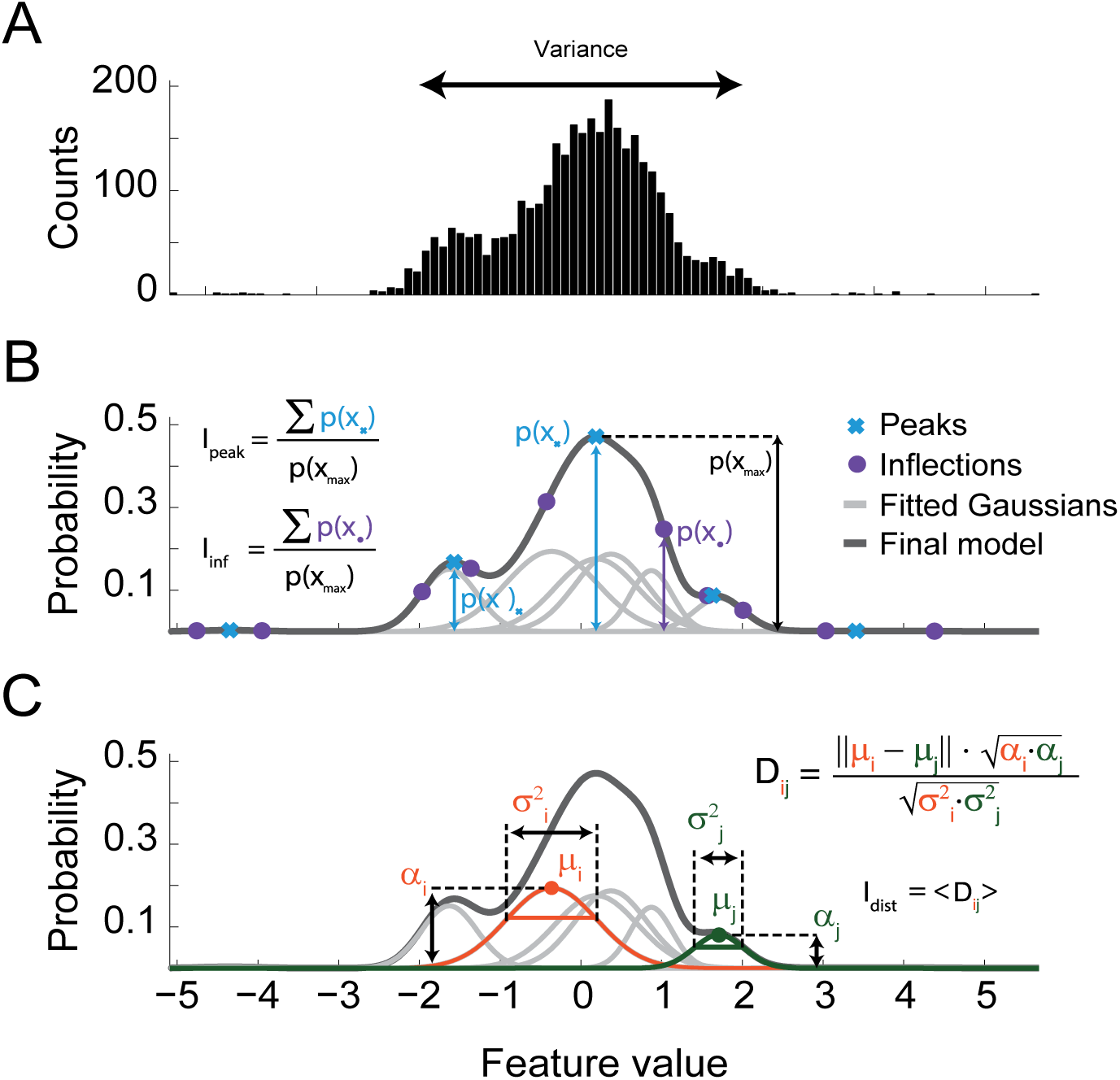
Estimating clustering information of a feature with Gaussian mixture models. **A.** Histogram of the values of a waveform feature (e.g., PC scores or wavelet coefficients). Note multimodal distribution. The variance of the feature values (var) was one of the 4 separability metrics studied in this work. **B.** Gaussian mixture model of the feature values in A. The black line shows the probability distribution function of the mixed model composed by eight Gaussians (gray lines). Crosses and disks mark the peak and inflection points of the model, respectively, which were used to compute two of the separability metrics (I_peak_ and I_inf_). **C.** Schematic representation of the normalized distance between Gaussian pairs (D_ij_); μ: Gaussian mean; σ: standard deviation; α: Gaussian weight. Another separability metric was defined as the median distance (I_dist_).

The fourth metric we used was the variance of the features, which is independent of the GMM. For each of the four metrics, we selected the 5 highest-ranked PC scores or wavelet coefficients to perform clustering.

In addition to pure WD, we also implemented a third technique for feature extraction using wPCA in the wavelet coefficients. To that end, each coefficient was z-scored and multiplied by one of the three GMM-based metrics (I_peak_, I_inf_ or I_dist_) prior to computing PCA (notice that we did not use the variance metric since this would oppose the z-score normalization). The first 5 weighted PC scores (ranked by variance) were then selected for clustering (see S1 Fig).

#### Clustering

After feature extraction (5 features for each combination of technique and clustering metric), we first estimated the number of clusters in the data using an overclustering approach. To do so, we fitted a GMM for the data pro.ected into the 5 selected features with a large number of Gaussians (12 for Dataset A and 20 for Dataset B; see below) (Fig 3B,C). In this model, each Gaussian was defined by their mean (center) and covariance in the 5-feature space. We then searched for peaks in the GMM probability function surface by running a Nelder-Mead simplex algorithm [24] for 12 or 20 initial conditions, defined by the Gaussian centers. Peaks distant by less than our resolution (100 bins) were merged and counted as one. Finally, each peak of the mixture was then regarded as a cluster center (Fig 3C).

**Fig. 3.**
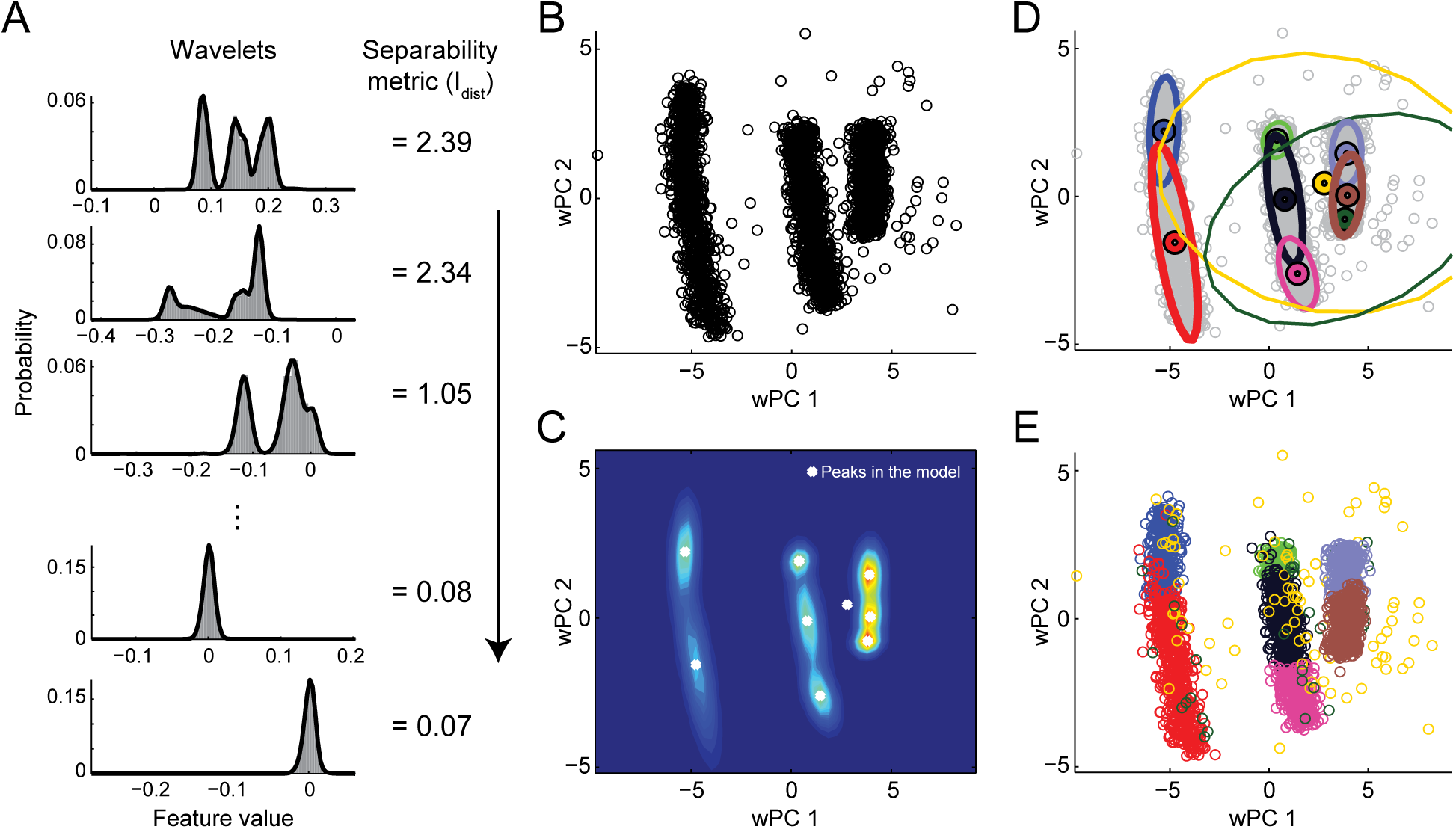
GMM-based spike sorting framework. **A.** Example of feature values (wavelet coefficients) and their GMM fittings. In the proposed framework, wavelet coefficients (or PC scores) are ranked according to a clustering separability metric; 4 metrics were investigated: var, I_peak_, I_inf_ or I_dist_ (see Materials and Methods). The example depicts wavelet coefficient distributions ranked by the I_dist_ metric. Note that unimodal distributions have lower I_dist_ values. **B.** The first two wPCs of the coefficients in A. Weighted PCA was obtained by normalizing the variance of wavelet coefficients by I_dist_ and applying PCA. Clustering was done using the first 5 wPCs. For pure PCA and WD, we used the first 5 features according to the separability metric. **C.** The probability density function of the 5-dimension GMM computed from the feature subspace (same dimensions as in B) and its peaks (white dots). The GMM was computed with 12 (Dataset A) or 20 (Dataset B) Gaussians. **D.** Representation of a fixed-mean GMM with Gaussians centered at the peaks in C. Dots and ellipses denote the center and the 2-standard-deviation boundary of each Gaussian; line thickness represents Gaussian amplitude. **E.** Final classification of the waveforms using the fixed-mean GMM. Each waveform was assigned to the Gaussian with higher probability in the model in D. Colors in D and E represent different Gaussians/clusters.

To classify the waveforms into different clusters, we performed yet another GMM fitting using the cluster centers found in the previous step as fixed means (i.e., a fixed-mean GMM in the 5-feature space). In this case, the number of Gaussians was equal to the number of cluster centers. The Gaussians in this model defined cluster probabilities in the feature space for clustering (Fig 3D). Thus, we assigned each 5-dimensional point (representing a waveform) to the cluster of highest a posteriori probability (Fig 3E).

#### Comparing sorting performance

We compared the different strategies (PCA, WD and wPCA) of our GMM approach within themselves (I_peak_, I_inf_, I_dist_ and variance), with each other, and with two other spike sorting methods based on mixture models: KlustaKwik [9] and EToS [11]. To this end, we analyzed two datasets available online that were used in previous spike sorting studies [8,14] (S1 Table).

Dataset A was composed of 20 simulated recordings, each containing three different neurons [14]. Briefly, background activity consisted of spikes randomly drawn from a database containing 594 average waveforms. Three waveforms from the same database were superimposed on the background activity at random times with different signal-to-noise ratios.

Dataset B was composed of simultaneous intra and extracellular recordings of CA1 pyramidal cells in anesthetized rats [8]. In this case, the intracellular recording served as a ground truth for the cellular identity of one of the extracellular spikes. We tested the spike sorting methods using tetrode recordings available in this dataset. A detailed description of datasets A and B can be found in references [14] and [8], respectively.

#### Measuring classification performance

We used the mutual information (MI) [25] between the real and assigned spike classes to measure classification performance (S2 Fig). MI is defined as:

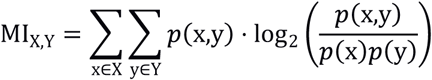

where X and Y are the real and assigned classes. The MI measures how mixed the real classes are within the assigned classes, providing the entropy shared by them. This is particularly valuable since unsupervised clustering can assign waveforms into a different number of clusters than the real one. S2 Fig B,C shows how the MI is affected by adding more assigned classes or mixing them. In order to compare the MI between different waveform sets, we normalized each MI by its maximum possible value (i.e., the entropy of the real class). This yields the percentage of extracted spike information (MI_norm_):

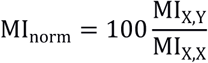

We computed MIs through the Information Breakdown toolbox [26].

For Dataset A, we subsampled spikes to investigate the effect of unbalanced clusters (different firing rates) on classification performance. More specifically, we initially set the three neurons to have the same number of spikes; we next subsampled the waveforms of one of the neurons to a percentage of its total number of spikes, which defined the “symmetry index” (i.e., a symmetry index of 10% corresponds to a reduction to 10% of the total number of spikes). Each of the three neurons had their spikes subsampled individually, with symmetry indexes varying logarithmically from 1 to 100%. For this analysis, we also computed the MI_norm_ using the mutual information between the classification of the subsampled neuron against the others (MI_norm_ of the smallest cluster). All analyses were implemented in MATLAB.

## Results

We first used simulated data (Dataset A) to investigate how well the spike sorting approaches separate the neurons. To that end, we used the mean percentage of extracted spike information (mean MI_norm_) over the multiple runs of each set of waveforms (Fig 4A). We found that PCA-based strategies had variable performance regardless of the information metric used, and presented a broad distribution of mean MI_norm_. On the other hand, WD and wPCA had mean MI_norm_ values concentrated around 90%, except when using WD with variance, whose distribution was similar to PCA-based strategies. We found that EToS also had a good performance (~90% of mean extracted spike information), while KlustaKwik had lower performance (~60%).

**Fig. 4.**
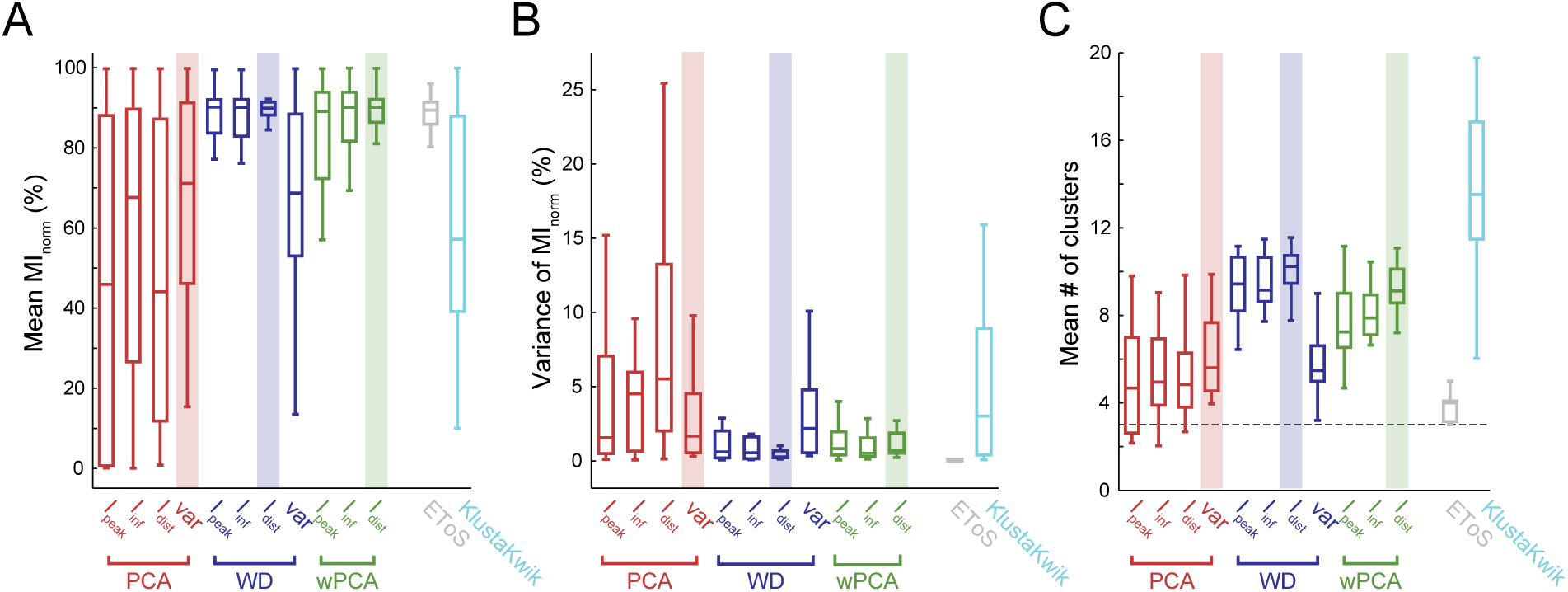
Spike sorting performance in a simulated dataset. **A.** Boxplots of mean spike information (MI_norm_) for each GMM-based feature extraction strategy. Sorting was performed 25 times for 20 sets of 3 neurons from Dataset A. Mean MI_norm_ values for EToS and KlustaKwik are also shown. **B-C**. Boxplots of MI_norm_ variance (B) and mean number of detected clusters (C) across the 25 runs of each set of neurons. Note that since the GMM-based methods are based on overclustering, the number of clusters was always higher than the true number of neurons (dashed line). Merging of clusters is a common post-processing step for several sorters. Highlights show the best (high performance and low variance) metric of cluster information for each feature extraction approach. The var metric was the most informative for PCA (red), while I_dist_ was the best metric for WD (blue) and wPCA (green).

We next investigated the consistency of the sorting results across runs, defined as the variance of MI_norm_. In other words, this measures if the algorithm provides similar MI_norm_ values across multiple runs of the same data. The PCA-based strategies and the WD-variance case had lower consistency (high variance of MI_norm_) than wPCA and the other WD strategies in simulated data (Fig 4B). EToS showed almost no variance, while KlustaKwik had low consistency across runs. We also investigated the mean number of clusters found in each run. Among GMM methods, the PCA-based strategies and WD-variance had the lowest mean number of clusters, followed by wPCA and the other WD strategies (Fig 4C). EToS had the lowest number of clusters, while KlustaKwik the highest. Based on the mean and variance (consistency) of extracted spike information, we selected the variance metric for PCA and the I_dist_ metric for either WD or wPCA as the best strategies (highlighted in Fig 4).

We performed the same analysis for real neuron data recorded from tetrodes (Dataset B). To that end, we combined intracellularly confirmed single units from different recordings in groups of 5 or 10. We found that the mean extracted spike information (mean MI_norm_) decreased as the number of neurons increased (Fig 5A), and that the best strategies selected for the simulated dataset were also among the best in the real dataset (highlights in Fig 5). The best wPCA and PCA approaches had similar mean MI_norm_, slightly outperforming the best WD method. Surprisingly, KlustaKwik had higher performance than EToS. The consistency results (Fig 5B) and the mean number of clusters (Fig 5C) found with each method exhibited comparable relations among metrics as in simulated data (compare with Fig 4B,C). One exception, however, was that KlustaKwik exhibited the highest consistency.

**Fig. 5.**
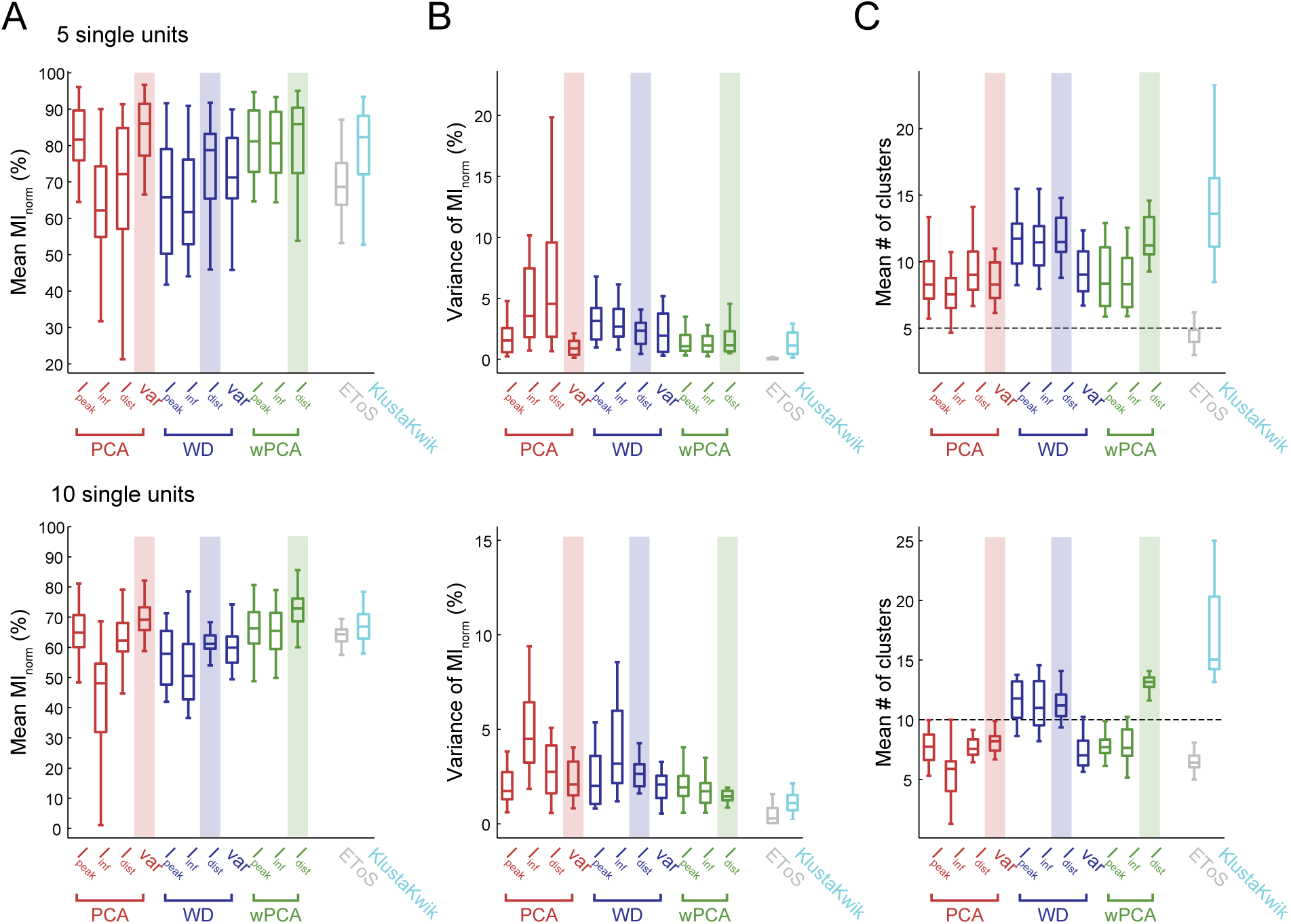
Spike sorting performance using real waveforms. Single units from Dataset B were pulled together in groups of 5 (top) or 10 (bottom) and used to test spike sorting performance. In this dataset, the identity of the extracellularly recorded waveforms was confirmed through simultaneous intracellular recordings. **A-C**. Mean spike information (MI_norm_) (A), MI_norm_ variance (B), and the mean number of detected clusters (C) are shown as in Fig 4.

We next compared the performance of the GMM-based best strategies (variance for PCA and I_dist_ for WD and wPCA) with EToS or KlustaKwik in a pairwise manner for Dataset A (Fig 6) and Dataset B (Fig 7). For Dataset A, we found that WD, wPCA and PCA outperformed KlustaKwik, and that WD and wPCA were similar to EToS (Fig 6). For Dataset B, we found that WD was similar to EToS while the PCA and wPCA strategies had mean MI_norm_ values higher than EToS and slightly better than KlustaKwik (Fig 7).

**Fig. 6.**
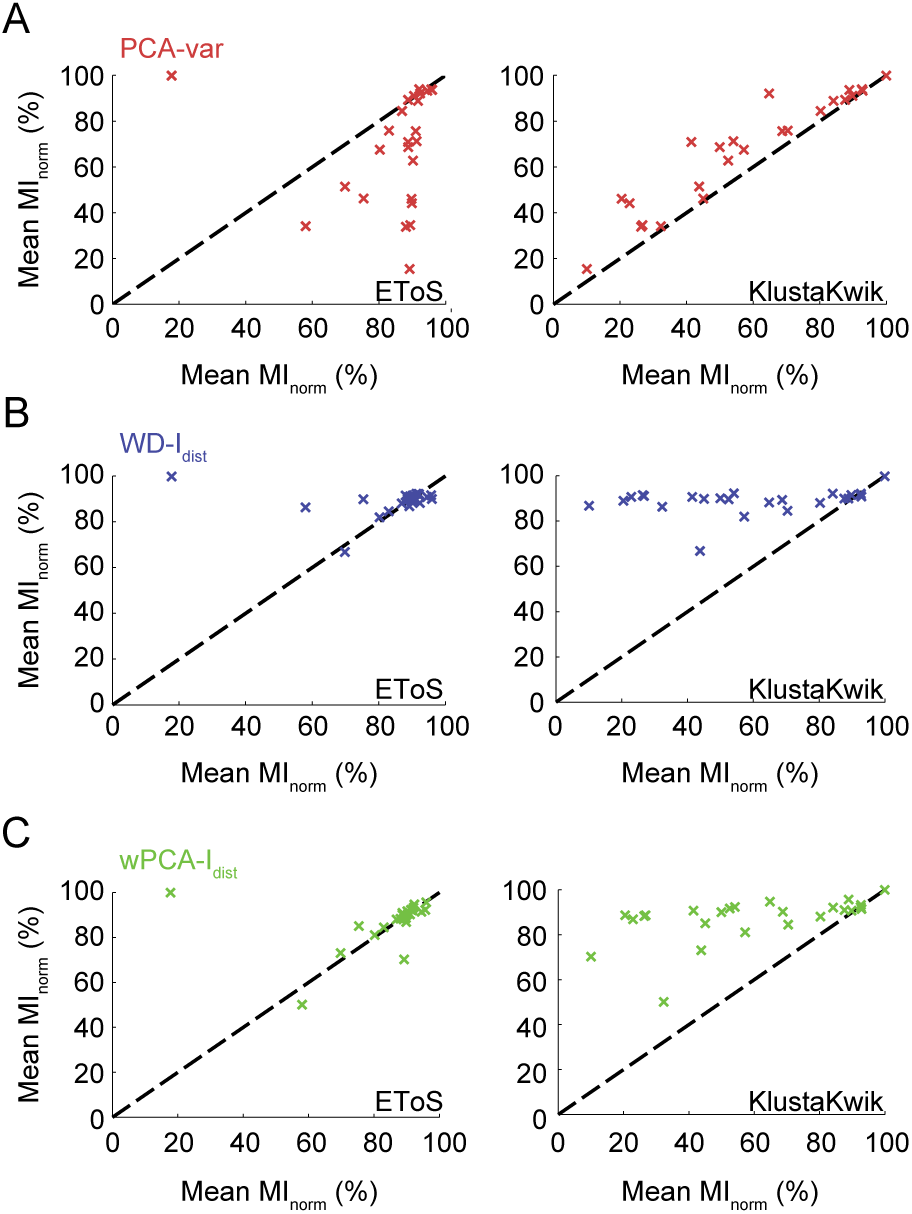
Comparing sorting performance in Dataset A. **A-C.** Pair-wise comparison between the best strategy for PCA (A), WD (B) or wPCA (C) and EToS or KlustaKwik. All strategies performed better than KlustaKwik, while WD-I_dist_ and wPCA‐‐I_dist_ performed similarly to EToS.

**Fig. 7.**
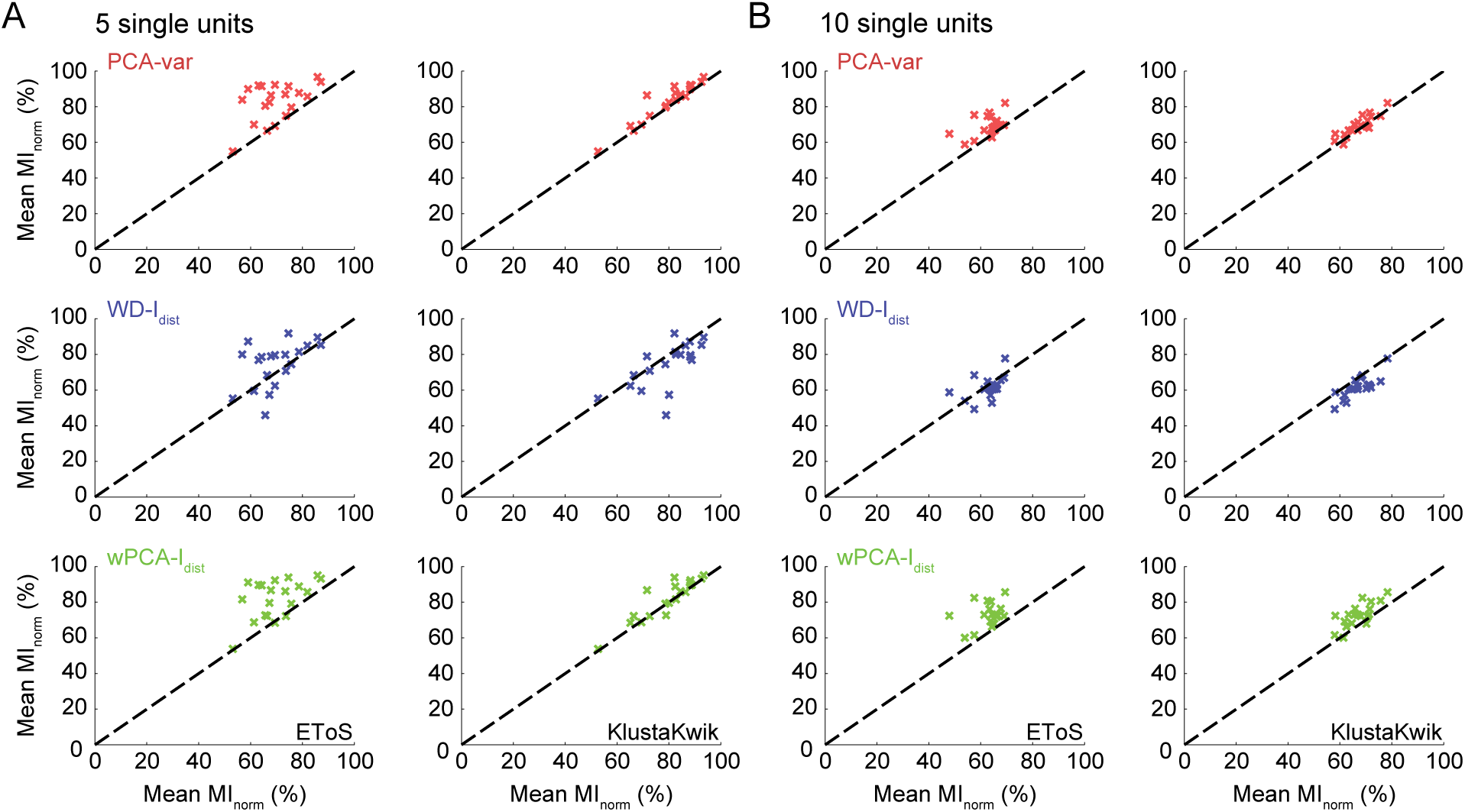
Comparing sorting performance in Dataset B. **A-B.** Pairwise comparison between the best PCA, WD or wPCA strategies and EToS or KlustaKwik for groups of 5 (A) and 10 (B) single units in Dataset B. PCA-var and wPCA-I_dist_ performed better than EToS and slightly better than KlustaKwik, while WD had similar performance to these methods.

We next investigated how each algorithm performs at the different signal-to-noise ratios present in Dataset A (Fig 8A). We computed the mean MI_norm_ and the number of clusters for each set of waveforms at varying degrees of noise levels (Fig 8B,C). The MI_norm_ values of both PCA-variance and KlustaKwik were the most sensitive to signal-to-noise ratio, rapidly decreasing as the noise level increased. Regarding the mean number of clusters, the EToS algorithm was the most robust across noise levels, and provided the best estimates for the number of clusters in the data. For the PCA, WD and wPCA strategies, the mean number of clusters tended to decrease with the noise level, whereas KlustaKwik exhibited the opposite trend.

**Fig. 8.**
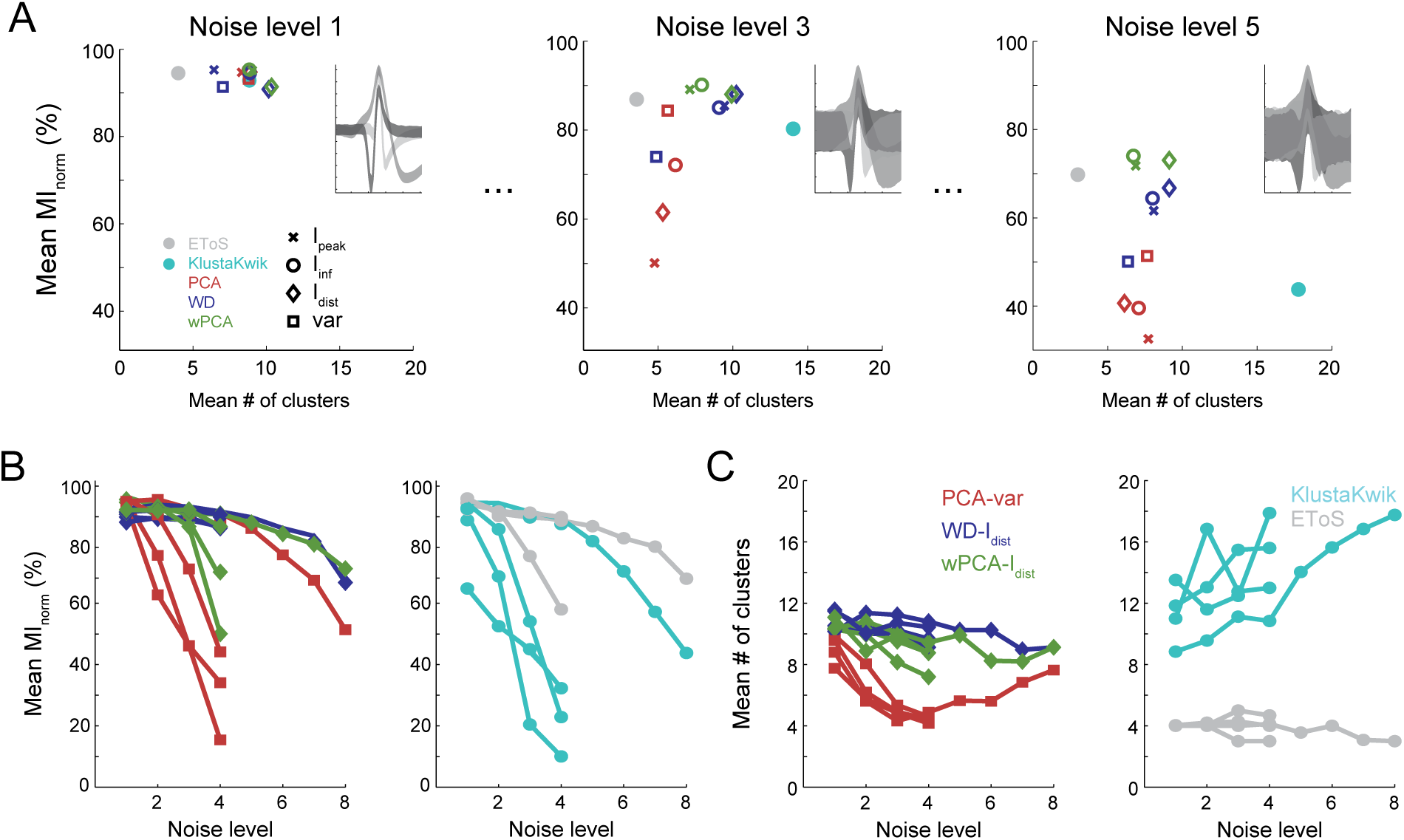
Spike sorting performance at different noise levels. **A.** Average spike information (MI_norm_) and number of detected clusters for the evaluated spike sorters in three example datasets (same set of 3 neurons differing in noise levels). The inset shows the 90% percent confidence interval (shadows) of the waveforms. **B-C.** Mean MI_norm_ (B) and number of clusters (C) for all datasets in Dataset A plotted as a function of the noise level. The lines connect datasets formed by the same set of 3 neurons. Note that as noise increases, the classification performance decreases.

Asymmetrical cluster sizes arise whenever neurons have very different firing rates (e.g., pyramidal cells and interneurons), which is often the case. We have thus analyzed the effects of unbalanced cluster sizes on clustering performance. To that end, we defined a symmetry index (see Materials and Methods), and randomly subsampled the datasets to match different values of symmetry index prior to performing the sorting procedure. We found that the mean MI_norm_ was relatively uncorrelated with the symmetry index (Fig 9A). However, when we computed a modified MI_norm_, which takes into account only the information extracted from the smallest cluster, we found that this metric decreased for lower symmetry indexes (Fig 9B), indicating that spikes from smaller clusters tend to be integrated into larger ones. In other words, the capacity of discriminating the smaller cluster lowers as the asymmetry in cluster size increases. WD and wPCA approaches were the least affected by unbalanced cluster sizes.

**Fig. 9.**
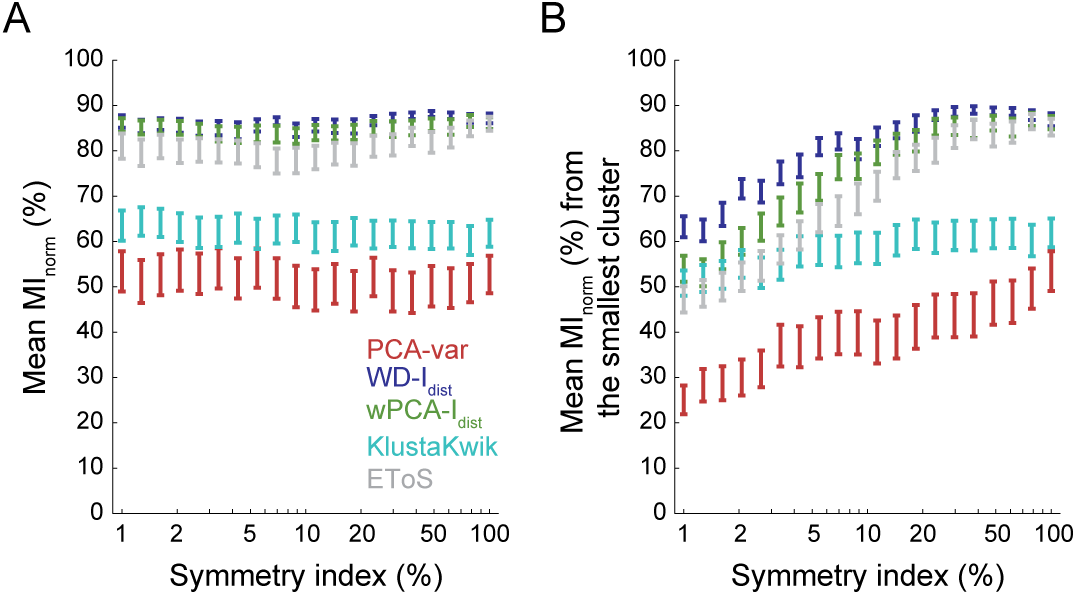
Sorting performance in unbalanced cluster sizes. **A.** Average spike information (MI_norm_) for sets of neurons with different cluster sizes (symmetry index). There were no apparent changes in MI_norm_ across different symmetry indexes. **B.** Similar to A, but showing only the spike information extracted from the smallest cluster. For this analysis, the other two clusters were joined prior to computing the MI_norm_ (see Materials and Methods). Despite the apparent constancy of the overall information (A), the classification of the smallest cluster decreases with the cluster size (B).

Based on the results above, we considered the GMM-based wPCA-I_dist_ algorithm as the one with best overall performance. We created a friendly graphical user interface to run the wPCA-I_dist_ algorithm and inspect the resulting classification (S3 Fig), which also allows for manual adjustments of cluster identity (e.g., merging of clusters). The codes and instructions are freely available at https://github.com/tortlab/GMM-spike-sorting.

## Discussion

In this work, we explored multiple strategies of feature extraction and selection using GMMs. We proposed GMM-based approaches to classify features and estimate the number of clusters in a data-driven way. These were compared to two spike sorting algorithms that are also based on mixture models [9,11].

The clustering step proposed here is based on an initial overclustering of the data (Fig 3). We first built a GMM of the selected features which overestimated the number of clusters, resulting in a mixture model with more Gaussians than the real number of neurons. Then, we estimated the cluster number – along with their centers – through the peaks of the mixed probability of the model. This is possible because adding new Gaussians decreases the error of the model. Thus, the few extra Gaussians (beyond the actual number of clusters) can be seen as a smoothing of the probability function of the mixed model that does not change the number and position of its peaks. Using the peak positions as new Gaussian centers, we recalculated the GMM and defined the cluster regions based on the new Gaussian distributions. These determined the maximum a posteriori probability used for clustering.

We used known feature extraction techniques (PCA, WD and wPCA) and combined them with three unsupervised estimators of clustering separability. The variance, which is widely used in standard PCA approaches, was used as a fourth clustering metric. We found that the variance was the best estimator of separability for the PCA-based approaches. This is in accordance to the standard use of the PCA in feature extraction, which selects the PC scores of highest variance [4,9,21]. However, for the simulated dataset (Dataset A), the PCA-variance performance was below WD and wPCA approaches, especially when combining them with the I_dist_ metric (Fig 4A). This supports previous work showing that WD outperforms PCA for spike sorting [13,14,27]. Nevertheless, the standard PCA presented better performance than WD in a second dataset composed of actual neurons (Dataset B; Fig 5A). Only wPCA, which combines both PCA and WD into a single approach, was among the best methods in both datasets.

We compared the performance of our sorting approaches to previously established algorithms: EToS and KlustaKwik, which are also based on mixture models [9,11]. Their performance varied according to the dataset used. KlustaKwik, which is based on PCA, performed better for Dataset B (Fig 5A), as it was the case of our own PCA approach. This might be explained by the fact that earlier versions of KlustaKwik were developed using Dataset B. On the other hand, EToS, which also uses a combination of wavelets and wPCA, was best suited for Dataset A (Fig 4A), similar to our WD approach. For each of the two datasets, our wPCA approach yielded better or comparable results to EToS and KlustaKwik, indicating it to be a good alternative for spike sorting (Figs 6 and 7).

We next investigated how each algorithm performs at the different noise levels in Dataset A. The WD approach had the most robust performance, and it was closely followed by wPCA and EToS. On the other hand, we found that PCA and KlustaKwik were more sensitive, and rapidly decreased their performance as the noise level increased (Fig 8B). These results support previous findings showing that the first PC scores capture a higher percentage of the noise energy when compared to WD [13].

Although we investigated sorting performance under different noise levels, we do not know how each method would perform in the presence of outliers. Due to the overclustering strategy employed in our approach, it is possible that outliers generate an extra cluster arising from a Gaussian with high standard deviation and low amplitude. Previous work suggested that, because of its longer tail, a mixture model of t-Student distributions (instead of Gaussians) would be more indicated to deal with outliers [21]. However, for Dataset B our method showed better performance than EToS, which is based on t-Student’s mixture model. Since they use different feature extraction methods, it is unknown whether changing Gaussians for t-distributions in our case would improve its performance. Nevertheless, our method can be adapted to other types of mixture models. In fact, the clustering and feature selection approaches proposed here can be used independently and combined with other strategies (e.g., our feature selection strategy could be combined with KlustaKwik, or our overclustering approach could be applied to other waveform features not investigated here).

For some clustering strategies, better performance was accompanied by an increased number of detected clusters (Figs 4C and 5C). Since high-performance approaches had high MI_norm_ values, this might be due to splitting the waveforms of a same neuron into more than one cluster (see S2 Fig and Fig 3E). This brings into question whether it is possible to optimize both classification performance (here defined as MI_norm_) and the estimation of the correct number of clusters by a sorting algorithm. Spike sorters use differences in waveform shapes to separate neurons. However, the spikes of a same neuron can eventually have different waveforms shapes (e.g., the later spikes in a burst), and the clustering step will not be able to distinguish these cases from spikes arising from different neurons. Therefore, unless the classifier is able to use other variables to aid in the assignment of neuronal identity (e.g., spike timing), posterior adjustments are often required, either by a manual operator or another processing algorithm. Of note, in our GMM-based framework, merging of clusters is currently done manually using the GUI we developed (S3 Fig). Notice that from a practical point of view it is simpler to merge clusters a posteriori than to separate them. In fact, a variety of spike sorting methods have been proposed in recent years with improved capabilities of neuronal classification [9,14,21,28], but still depend on parameter adjusting and manual curation steps [29]. Recent work has shown efforts for implementing a fully automated spike sorting [30], encapsulating these postprocessing steps within the algorithm. However, this approach entirely focuses on the clustering stage, which leaves unanswered the question of how much feature extraction techniques can improve the final outcome.

We also analyzed the performance of our approaches under different conditions of symmetry and found that the WD was the most robust algorithm, closely followed by wPCA and EToS. This is important because neurons with lower firing rates can go undetected, assimilated by larger clusters [31,32]. Notably, because each Gaussian in the mixture is fitted independently, GMMs (or other mixture models) offer a good solution to this problem as they can capture clusters of different size, assigning them to Gaussians with different standard deviations and weights. The GMM approach constitutes a free-scale smoothing whenever the data comes from multiple distributions with different scales (standard deviations), eliminating the need for defining the length (scale) of the smoothing function.

In summary, our results bring new tools to tackle the spike sorting problem. Concerning the initial steps of feature extraction, we proposed fitting a GMM to wavelet coefficients or PC scores to estimate its clustering separability under a variety of metrics. Our results show that the combination of WD and PCA (the wPCA) is a better approach than using one of them separately as employed by most spike sorters [9,13,14,30]. Namely, the wPCA associated with a simple metric of distance between Gaussians (I_dist_) was the best strategy of feature extraction in the two investigated datasets. Finally, in the step of unsupervised clustering, we proposed fitting an overclustered GMM, searching for local maxima to estimate cluster positions and then re-fitting the GMM with Gaussians at the estimated positions. This simple strategy can also be combined with multiples runs with different numbers of Gaussians, as done in previous studies [33,34]. We hope these findings shed new light on spike sorting and other unsupervised clustering problems.

## Supporting Information

**S1 Fig.**
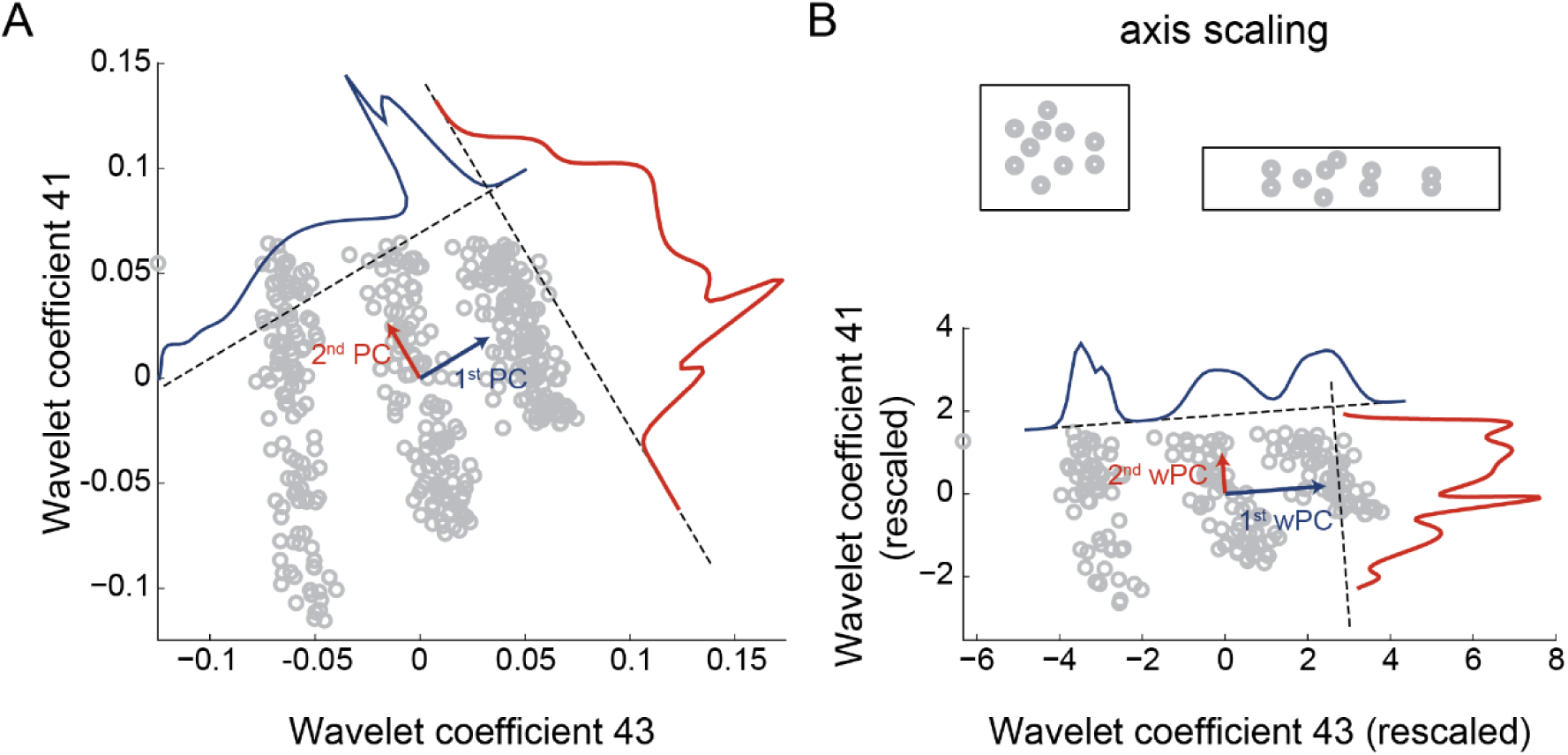
Rescaling wavelet coefficients by separability metric. **A.** Scatter plot of two wavelet coefficients. The first and second principal components (blue and red arrows) are shown along with a Gaussian fit of the data projected onto these axes. **B.** (Top) Scheme of axis rescaling. In the weighted-PCA each dimension is z-scored and scaled by a particular metric before the computation of the principal components (i.e., the variance of each dimension is weighted). (Bottom) The same as in A after weighting the coefficients by the separability metric. Note that the first weighted principal component (blue curve in B) can better separate the three clusters than the first principal component (blue curve in A).

**S2 Fig.**
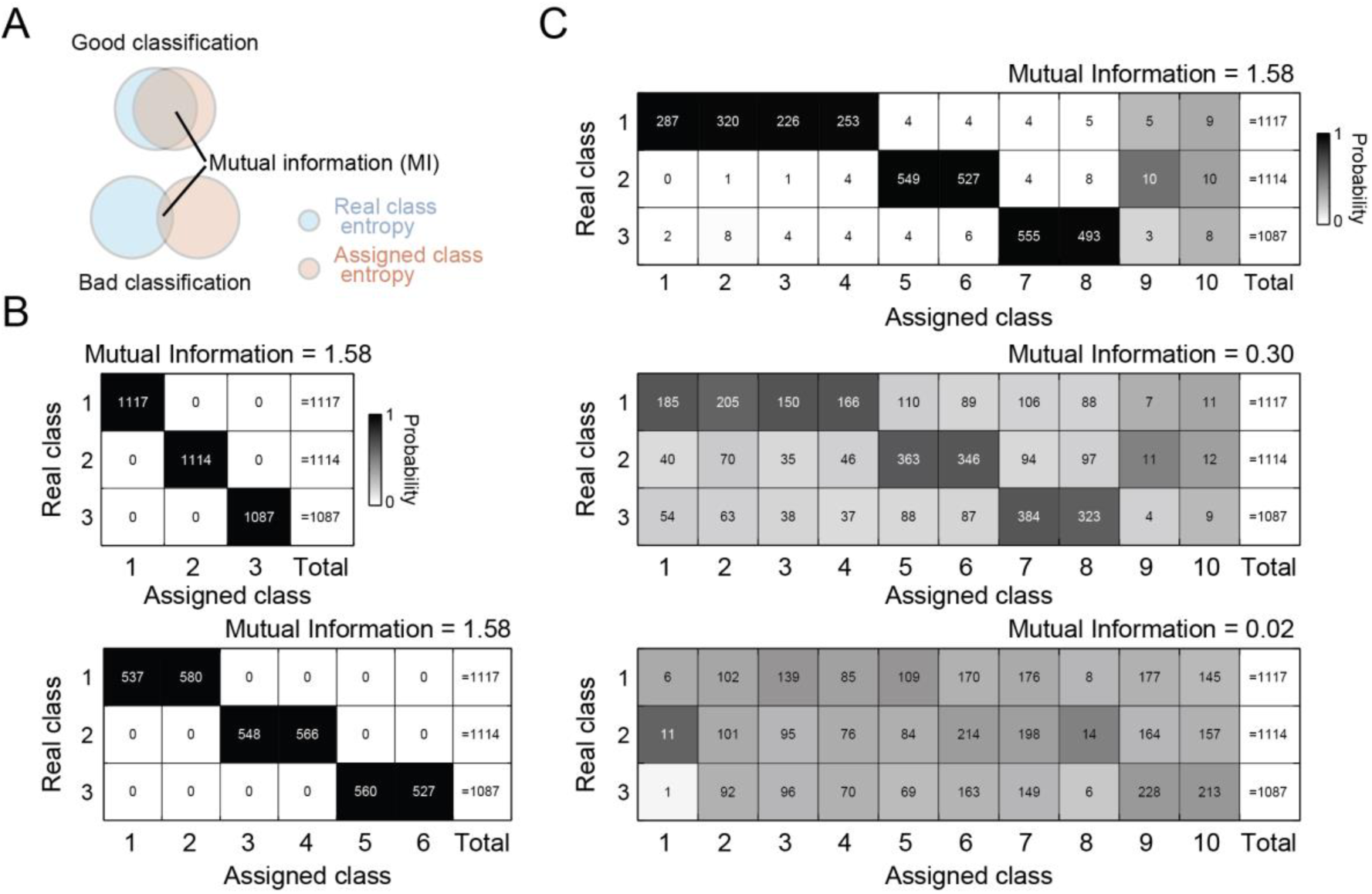
Measuring classification performance with Mutual Information (MI). **A.** Information diagram of real and assigned classes (in our case, neuronal clusters). Good performance implies higher mutual information (MI) between the two groups. **B.** Contingency tables showing perfect classification without (top) and with (bottom) overclustering. Table color denotes the probability (over columns) of each real class to be classified in that particular cluster (assigned class). **C.** Contingency table and MI values for 3 sorting examples differing in classification performance. Note that although the MI values are the same for both cases in B, its value is sensitive to the mixing of classes in C. The MI_norm_ shown in the main figures is defined as the normalization of the measured MI value by the maximal possible MI.

**S3 Fig.**
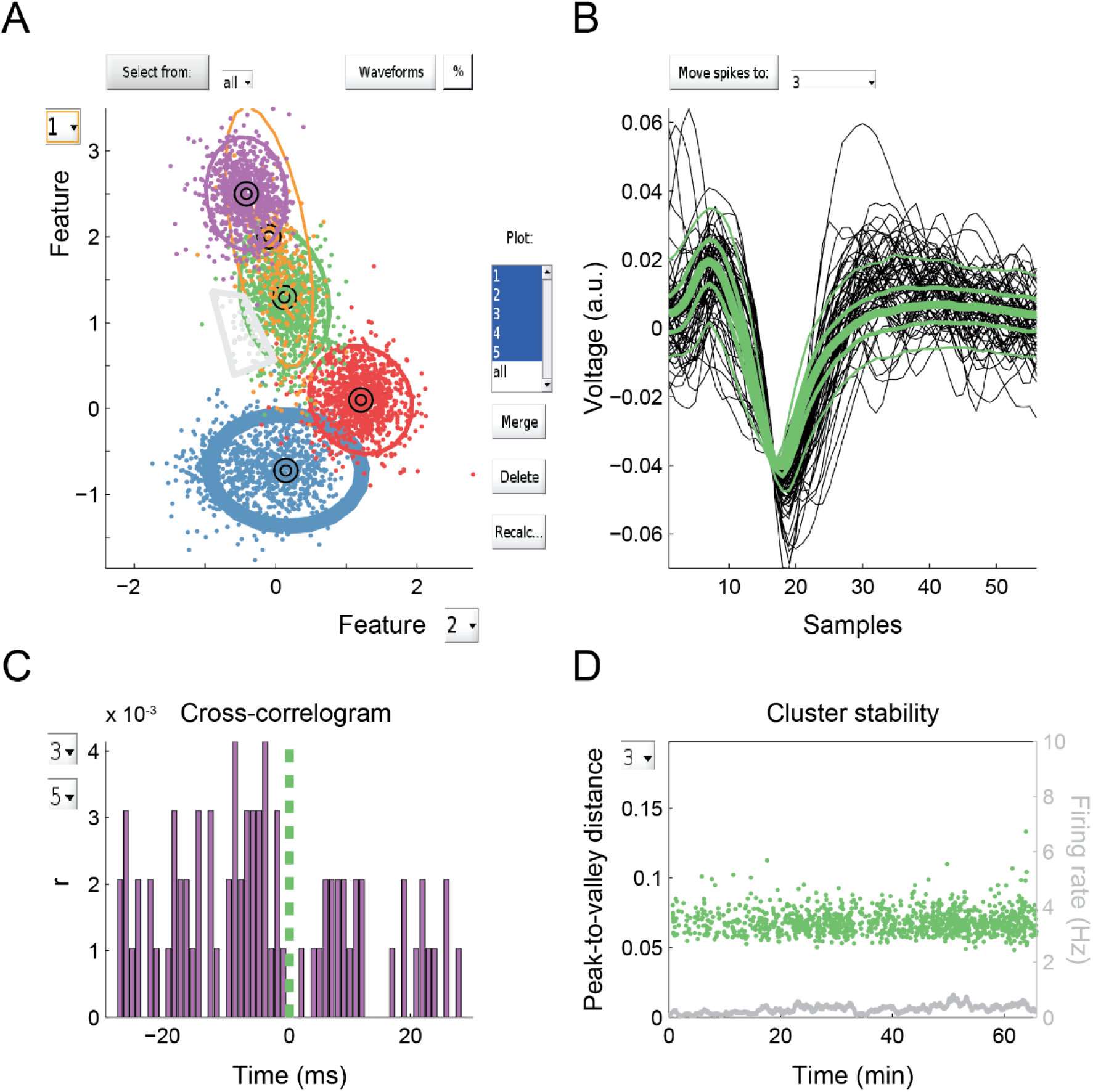
Example of a screen of the developed GUI. **A.** Feature space showing the first two out of five features. Ellipses demark Gaussians defining clusters. Gray polygon was selected manually for further inspection in B. **B.** Mean waveform for cluster 3 (green; thin lines show quantiles). Black waveforms correspond to the gray dots selected in A. **C.** Cross-correlogram between clusters 3 and 5. **D.** Peak-to-valley distance for individual waveforms and firing rate showing the stability of cluster 3 along the recording time. MATLAB scripts and GUI instructions are available at https://github.com/tortlab/GMM-spike-sorting.

**S1 Table.** List of datasets used.

|  | File |
| --- | --- |
| Dataset A (Quiroga et al., 2004) | C_Difficult1_noise01 |
|  | C_Difficult1_noise02 |
|  | C_Difficult1_noise005 |
|  | C_Difficult1_noise015 |
|  | C_Difficult2_noise01 |
|  | C_Difficult2_noise02 |
|  | C_Difficult2_noise005 |
|  | C_Difficult2_noise015 |
|  | C_Easy1_noise01 |
|  | C_Easy1_noise02 |
|  | C_Easy1_noise03 |
|  | C_Easy1_noise04 |
|  | C_Easy1_noise005 |
|  | C_Easy1_noise015 |
|  | C_Easy1_noise025 |
|  | C_Easy1_noise035 |
|  | C_Easy2_noise01 |
|  | C_Easy2_noise02 |
|  | C_Easy2_noise005 |
|  | C_Easy2_noise015 |
| Dataset B (CRCNS: hc1) | d5331 |
|  | d5611 |
|  | d7212 |
|  | d11222 |
|  | d12821 |
|  | d14521 |
|  | d14531 |
|  | d15121 |
|  | d17013 |
|  | d17111 |
|  | d18711 |

## References

1. Csicsvari J, Hirase H, Czurkó A, Mamiya A, Buzsáki G. Oscillatory coupling of hippocampal pyramidal cells and interneurons in the behaving rat. J Neurosci. 1999;19: 274–287.

2. Ranck JB. Studies on single neurons in dorsal hippocampal formation and septum in unrestrained rats. Exp Neurol. 1973;41: 462–531.

3. Barthó P, Hirase H, Monconduit L, Zugaro M, Harris KD, Buzsáki G. Characterization of neocortical principal cells and interneurons by network interactions and extracellular features. J Neurophysiol. 2004;92: 600–608.

4. Lewicki MS. A review of methods for spike sorting: the detection and classification of neural action potentials. Netw Comput Neural Syst. 1998;9: R53–R78.

5. Brown EN, Kass RE, Mitra PP. Multiple neural spike train data analysis: state-of-the-art and future challenges. Nat Neurosci. 2004;7: 456–461.

6. Buzsáki G. Large-scale recording of neuronal ensembles. Nat Neurosci. 2004;7: 446–451.

7. Quiroga RQ. Spike sorting. Curr Biol. 2012;22: R45–R46.

8. Harris KD, Henze DA, Csicsvari J, Hirase H, Buzsáki G. Accuracy of tetrode spike separation as determined by simultaneous intracellular and extracellular measurements. J Neurophysiol. 2000;84: 401–414.

9. Kadir SN, Goodman DFM, Harris KD. High-dimensional cluster analysis with the masked EM algorithm. Neural Comput. 2014;26: 2379–2394.

10. Takekawa T, Isomura Y, Fukai T. Accurate spike sorting for multi-unit recordings. Eur J Neurosci. 2010;31: 263–272.

11. Takekawa T, Isomura Y, Fukai T. Spike sorting of heterogeneous neuron types by multimodality-weighted PCA and explicit robust variational Bayes. Front Neuroinformatics. 2012;6.

12. Wang H-Y, Wu X-J. Weighted PCA space and its application in face recognition. Machine Learning and Cybernetics, 2005 Proceedings of 2005 International Conference on. IEEE; 2005. pp. 4522–4527.

13. Hulata E, Segev R, Ben-Jacob E. A method for spike sorting and detection based on wavelet packets and Shannon’s mutual information. J Neurosci Methods. 2002;117: 1–12.

14. Quiroga R, Nadasdy Z, Ben-Shaul Y. Unsupervised spike detection and sorting with wavelets and superparamagnetic clustering. Neural Comput. 2004;16: 1661–1687.

15. Zouridakis G, Tam DC. Multi-unit spike discrimination using wavelet transforms. Comput Biol Med. 1997;27: 9–18.

16. Strang G, Nguyen T. Wavelets and filter banks. SIAM; 1996.

17. Jain AK, Duin RPW, Mao J. Statistical pattern recognition: A review. IEEE Trans Pattern Anal Mach Intell. 2000;22: 4–37.

18. Lewicki MS. Bayesian modeling and classification of neural signals. Neural Comput. 1994;6: 1005–1030.

19. Sahani M, Pezaris JS, Andersen RA. On the separation of signals from neighboring cells in tetrode recordings. Adv Neural Inf Process Syst. 1998; 222–228.

20. Peel D, McLachlan GJ. Robust mixture modelling using the t distribution. Stat Comput. 2000;10: 339–348.

21. Shoham S, Fellows MR, Normann RA. Robust, automatic spike sorting using mixtures of multivariate t-distributions. J Neurosci Methods. 2003;127: 111–122.

22. Mallat SG. A theory for multiresolution signal decomposition: the wavelet representation. IEEE Trans Pattern Anal Mach Intell. 1989;11: 674–693.

23. McLachlan G, Peel D. Finite mixture models. John Wiley & Sons; 2004.

24. Lagarias JC, Reeds JA, Wright MH, Wright PE. Convergence properties of the Nelder-Mead simplex method in low dimensions. SIAM J Optim. 1998;9: 112–147.

25. Shannon CE. A mathematical theory of communication, Part I, Part II. Bell Syst Tech J. 1948;27: 623–656.

26. Magri C, Whittingstall K, Singh V, Logothetis NK, Panzeri S. A toolbox for the fast information analysis of multiple-site LFP, EEG and spike train recordings. BMC Neurosci. 2009;10: 81.

27. Pavlov A, Makarov VA, Makarova I, Panetsos F. Sorting of neural spikes: When wavelet based methods outperform principal component analysis. Nat Comput. 2007;6: 269–281.

28. Rossant C, Kadir SN, Goodman DFM, Schulman J, Hunter MLD, Saleem AB, et al. Spike sorting for large, dense electrode arrays. Nat Neurosci. 2016;19: 634–641.

29. Hazan L, Zugaro M, Buzsáki G. Klusters, NeuroScope, NDManager: A free software suite for neurophysiological data processing and visualization. J Neurosci Methods. 2006;155: 207–216.

30. Chung JE, Magland JF, Barnett AH, Tolosa VM, Tooker AC, Lee KY, et al. A Fully automated approach to spike sorting. Neuron. 2017;95: 1381–1394.e6.

31. Quiroga RQ. Concept cells: the building blocks of declarative memory functions. Nat Rev Neurosci. 2012;13: 587–597.

32. Rey HG, Pedreira C, Quian Quiroga R. Past, present and future of spike sorting techniques. Brain Res Bull. 2015;119: 106–117.

33. Yamaguchi Y, Aota Y, McNaughton BL, Lipa P. Bimodality of theta phase precession in hippocampal place cells in freely running rats. J Neurophysiol. 2002;87: 2629–2642.

34. Zheng C, Bieri KW, Trettel SG, Colgin LL. The relationship between gamma frequency and running speed differs for slow and fast gamma rhythms in freely behaving rats: slow and fast gamma correlations with speed. Hippocampus. 2015;25: 924–938.

35. Diehl, G.W, Hon, O.J., Leutgeb, S., Leutgeb, J.K., 2017. Grid and Nongrid Cells in Medial Entorhinal Cortex Represent Spatial Location and Environmental Features with Complementary Coding Schemes. Neuron 94, 83–92.

